# Local low frequency fMRI synchrony is a proxy for the spatial extent of functional connectivity

**DOI:** 10.1101/138966

**Authors:** Gregory R. Kirk, Daniela Dentico, Rasmus Birn, Nagesh Adluru, Thomas Blumensath, Bill Taylor, Lauren Michael, Manuel F. Casanova, Andrew L. Alexander

**Author notes:** Corresponding Author: Gregory Kirk, Waisman Laboratory for Brain Imaging and Behavior, University of Wisconsin-Madison, 1500 Highland Avenue, Madison, WI 53705, USA.

## Abstract

Functional connectivity Magnetic Resonance Imaging (fcMRI) has assumed a central role in neuroimaging efforts to understand the changes underlying brain disorders. Current models of the spatial and temporal structure of fcMRI based connectivity contain strong a priori assumptions. We report that low temporal frequency fMRI signal synchrony within the local (3 mm radius) neighborhood of a location on the cortical surface strongly predicts the scale of its global functional connectivity. This relationship is tested vertex-wise across the cortex using Spearman’s rank order correlation on an individual subject basis. Furthermore, this relationship is shown to be dynamically preserved across repeated within session scans. These results provide a model free data driven method to visualize and quantitatively analyze patterns of connectivity at the imaging voxel resolution across the entire cortex on an individual subject basis. The procedure thus provides a tool to check directly the validity of spatial and temporal prior assumptions incorporated in the analysis of fcMRI data.

---

Since the first report of correlated low frequency blood oxygenation level dependent (BOLD) fMRI time series within functional areas^1^ resting state functional connectivity MRI (fcMRI) has evolved into a tool used to study distributed networks, termed functional connectivity networks, throughout the brain ^2, 3^. These low temporal frequency BOLD correlations have been shown to be related to neuronal activity patterns through optical imaging ^4^, electrophysiology ^5^, axonal tract tracing ^6^, tractography with diffusion-weighted imaging ^7^, the abolishment of correlations by surgical disconnection ^8, 9^ correlations with FDG-PET glucose measurements ^10^ and Transcranial Magnetic Stimulation ^11^. It is currently unknown whether fcMRI connectivity measurements reflect co-activation, the number of axons connecting the areas, strength of synaptic connections or dynamic connectivity modulated by the dozens of neuromodulatory chemical signals that are known to change neuronal connectivity on the cellular level^12^. Most likely the connectivity is constrained by the axonal and synaptic connectivity on long time scales and gated and modulated by the neuromodulatory influences on faster time scales^13, 14^.

Notwithstanding the uncertain biological interpretation of connectivity as measured by fcMRI, large efforts are underway to understand the spatial structure of connectivity throughout the brain^15, 16^. Functional area parcellations and the network analyses which use node structures derived from these parcellations decompose the brain or sub-regions of it into subdivisions which are by some measure of functionally connectivity differentiated. Many procedures have been described to perform this subdivision, most prominently seed clustering approaches ^6, 17, 18^ and subspace projection methods typified by Independent Component Analysis^19^. All of these are primarily data reduction procedures. They typically operate at a relatively large spatial scale and yet there is evidence of much more detailed structure^20^. These data reduction strategies can be compromised by deviations from the modeling assumptions used. One modeling assumption is related to temporal variation in functional connectivity. There is evidence that the degree of temporal dynamics has been underestimated and the influence of temporal changes in connectivity not sufficiently integrated into the analysis and interpretation of fcMRI studies^21, 22^. Group analyses may be affected by the additional unknown individual variability in structural and functional anatomy across subjects^23–26^.

The question arises as to the fidelity of the approximations afforded by these data reduction strategies. In graph analysis of functional connectivity, the brain or cortex is generally subdivided into some number of regions, or ‘nodes’, and either the average of all time series within the node or a chosen voxel or vertex within the node is used as a seed time series for seed correlation based approaches. For a given node definition, to what degree does the time series at an arbitrary or average seed within the node represent the time series of all other individual locations within the node. For subspace projection methods, to what degree does the properly weighted superposition of projected time series corresponding to the spatial components overlapping at a given point, reproduce the time series at that point, for all points across the cortex or brain gray matter volume.

Critically, very few analyses have been presented that assess the connectivity of every voxel or vertex independently across the entire cortex on an individual subject basis. Parcellations and graph models of the entire cortex have at most sampled several thousand locations across the cortex^27^. The at least implicit assumption in these approaches is that the cortex is approximately locally homogenous. Node definition is critical for valid network analyses. As elegantly described by Fornito and colleagues, “An ideal node definition for the human connectome should, define functionally homogenous nodes, represent functional heterogeneity across nodes, and account for spatial relationships” ^27^. The common practice in seed based fcMRI studies of using Talairach or MNI coordinates, defined from separate populations, to specify posited resting state network nodes implies a belief that the spatial homogeneity of resting state networks is high enough to accommodate the large and unquantifiable uncertainty in spatial registration using these methods ^24^. For graph analyses, predefined node definitions are often mapped onto the individual subjects in native space or normalized onto a template space. For analyses using large numbers of nodes, hundreds or thousands, low dimensional parcellations are either randomly or uniformly subdivided^27^. The random or uniform subdivision procedures are often not guided by any information regarding the actual connectivity of the subjects in the study. Therefore, it would be useful to have a procedure that indicates the spatial scale of connectivity of every functional voxel or vertex independently across the entire cortical surface on an individual subject basis. Such a procedure could be used to check the validity of a parcellation or node definition against actual connectivity at the individual subject level. A change in the spatial scale of connectivity between two points within a posited parcellation unit or network node would indicate that the connectivity is different between the two points and thus the parcellation unit or node is not functionally homogenous. Also importantly the same measure could be used to assess the degree to which the node boundaries coincide across subjects. In this study, we report evidence of a relationship between the synchrony of the local low temporal frequency BOLD signal within a 3 mm radius neighborhood of a vertex on the cortical surface and the spatial extent of the global functional connectivity of that vertex. The proposed analysis procedure is model free assuming no form of spatial prior and also contains no parameters that need to be chosen. The calculations performed are also very simple and so easily interpretable, but provide a wealth of information on the spatial structure of fcMRI connectivity.

## Results

The currently proposed measure is based on assessing the synchrony among the time series within a region of interest (ROI) as the mean square error between all time series within the ROI and the ROI’s mean time series. The ROI is here the set of all vertices within 3 mm of a target vertex on the cortical surface, the time series data having been resampled onto the individual subjects’ cortical surface model after preprocessing. The cortical surface was constructed using the Freesurfer software, an established and accurate method that performs cortical surface reconstruction based on the individual subjects structural T1-weighted MRI image data. Thus all locations on the cortical surface are referred to as vertices, a vertex being an element of the surface mesh used to model the 2-dimensional cortical surface. The synchrony measure is here termed Local Asynchrony (*LASY*) since larger values of the measure indicate more asynchronous time series and 0 indicates perfect synchrony. *We* also compute the seed based connectivity map of each vertex as the Pearson linear correlation between the time series of the vertex in question and the time series associated with all other vertices across both hemispheres of the cortical surface. We then threshold the resulting statistical parametric map and count the number of vertices with correlations above a specified reference level, thus achieving a measure of the connectivity extent (*CE*) associated with that vertex. Having obtained both these measures for every vertex across the cortical surface we are in a position to assess the relationship between *LASY* and CE. Note the chosen thresholds are only sampled to establish the existence of the inverse relationship between LASY and CE and are not needed to use the procedure once it has been established.

### The inverse relationship between *LASY* and *CE* is evident on a per scan basis

The data were preprocessed by motion correction, slice timing correction, and were band-pass filtered [0.1 − 0.01 hz]. The data were additionally processed to remove the mean white matter and CSF signals. Cardiac and respiratory signals collected at scan time were used to model and remove these influences (see Methods and supplementary material). The data were then projected onto the cortical surface and finally the time series were normalized to have zero mean and unit variance. Figure 1 presents an example of the empirical observation that framed our hypothesis of a relationship between *LASY* and CE. The top left and right figures show the cortex overlaid with the *LASY* measures resulting from two functional fMRI scans within the same scan session. These figures indicate the placement of seeds at the same vertex on the subject’s cortical surface. The corresponding correlation maps demonstrate the effect of using the fMRI time series at the seed vertex to perform seed based correlations analysis. The seed point on the left indicates high synchrony (low LASY) at the seed vertex and corresponds to a larger CE when compared to the connectivity map generated from the second resting fMRI data set now with the same seed vertex in a more asynchronous state (higher LASY).

**Figure 1.**
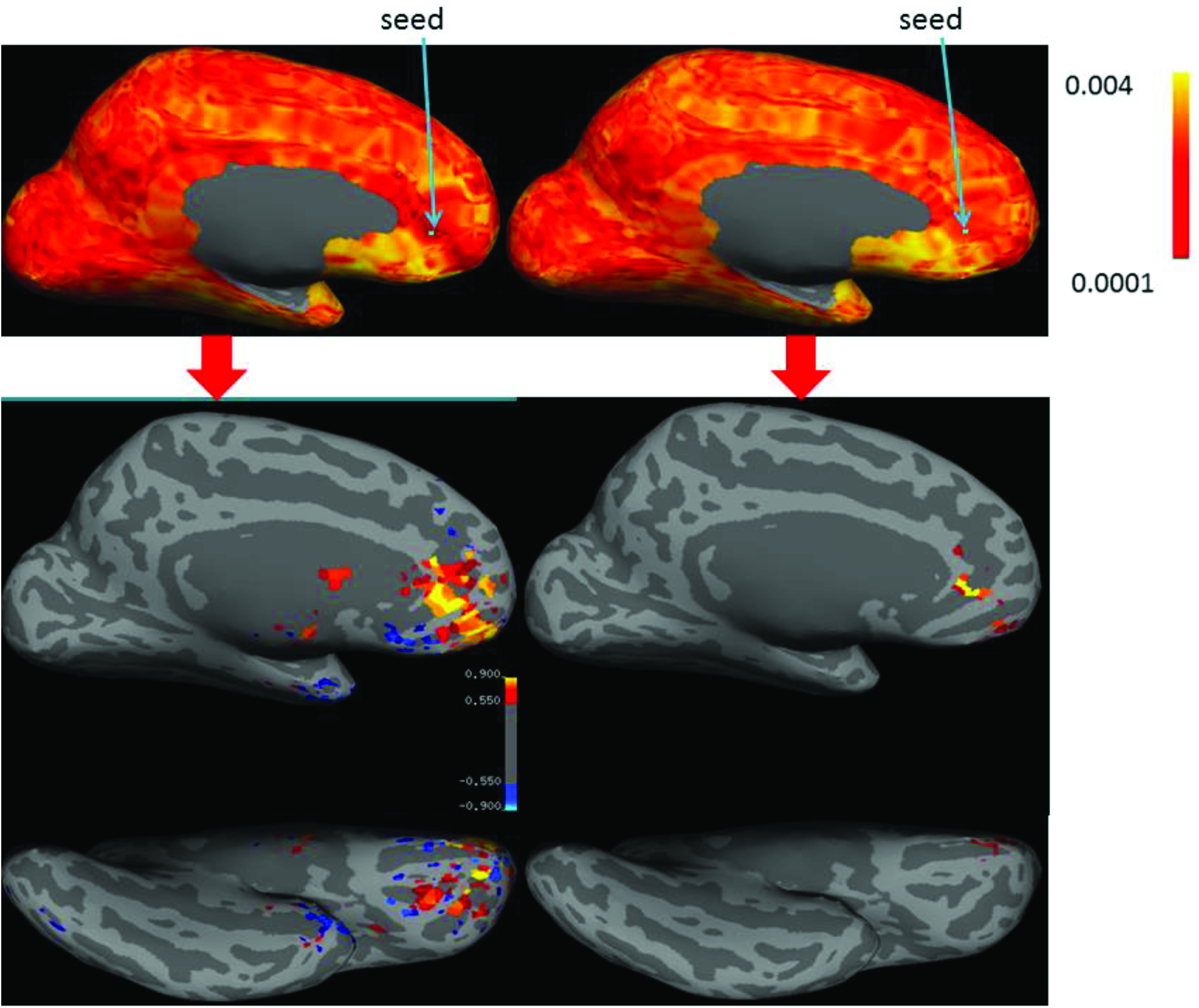
High synchrony implies larger networks. Top row displays *LASY* overlay calculated from two eyes closed scans from the same subject acquired during the same scan session. In both *LASY* overlay maps the blue arrow indicates the same vertex chosen as a seed location for Pearson linear correlation maps (indicated by red arrow) between the time series located at the seed and all other time series from the respective resting fMRI scans across the left medial and ventral aspect of the cortical surface. The correlation maps are thresholded at correlation coefficient > 0.55; vertices in yellow have correlations coefficients > 0.9. medial surfaces middle row. ventral surface bottom row.

The empirical observation that using vertices with synchronous local neighborhood as seed locations result in relatively large correlation maps, while vertices having an asynchronous local neighborhood correspond to smaller correlation maps forms the basis of our main hypothesis. The result is first demonstrated with an intuitive graphical representation of the data from a single scan.

Given there are approximately 300,000 vertices on both hemispheres of a subject’s cortical surface a direct plot of the relationship between *LASY and CE* is not practical. Therefore the *CE and LASY* data from a scan are partitioned into 100 element segments along the *LASY* axis. For each segment we calculate the mean *CE* value within the corresponding *CE* segment. This is depicted for an arbitrary 1000 element continuous segment at the top of figure 2. The red dots display the mean of the 100 individual vertex *CE* values (blue dots) within the segment.

**Figure 2.**
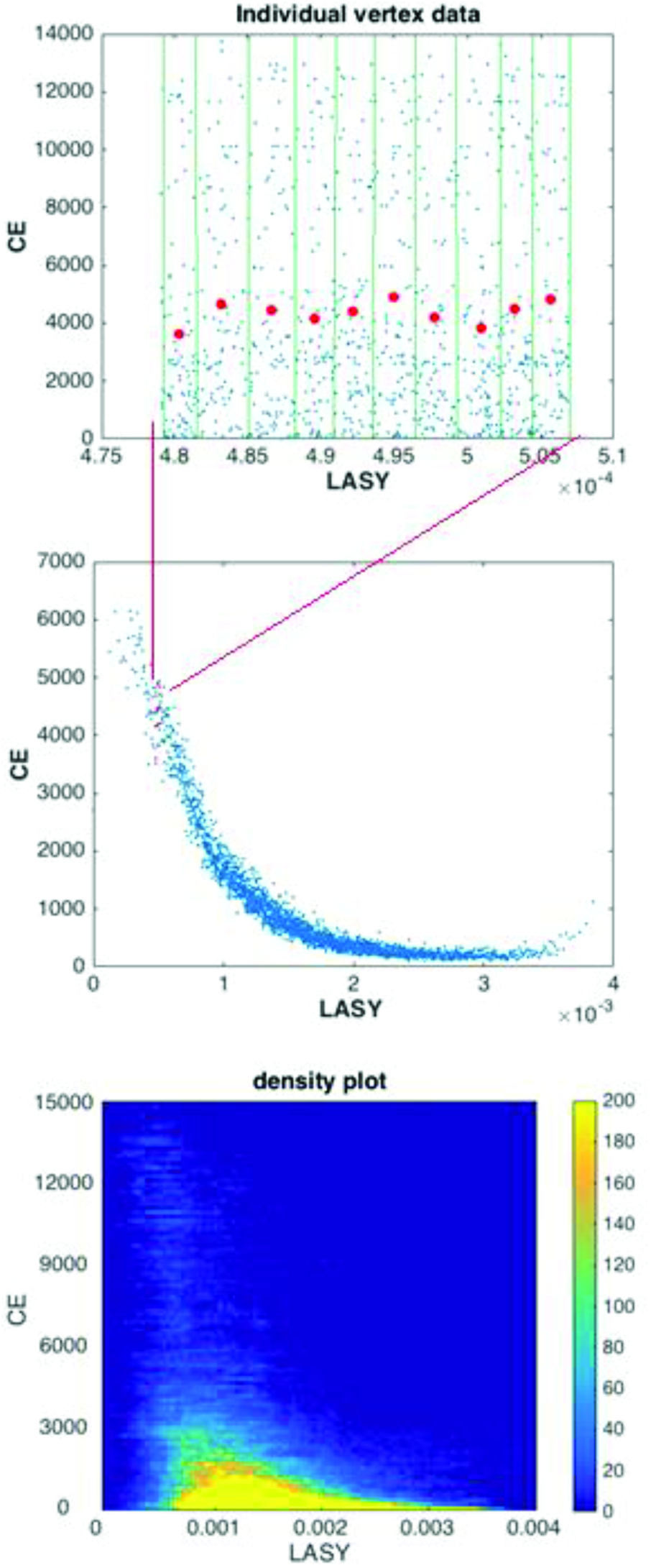
graphical representation of relationship between LASY and CE. A data reduction strategy is used in order to make interpretable the trend in the approximately 300,000 data point pairs contained in the *CE, LASY* data for a scan. In figure 2 top and middle row the data are split into non-overlapping 100 element segments on the *LASY* axis and the mean of the *CE* values is computed for each segment. This is depicted for a representative contiguous subset of 1000 data points where the individual 100 element segments are represented by the green lines in the top figure. The mean of each 100 element segments is indicated in red. The middle figure displays the mean of each 100 element segment for the entire data set as blue dots. The position from which the subset of 1000 data points was taken is indicated by the converging red lines. Figure 2 bottom provides the density plot of all ~300,000 pairs of the *CE, LASY* data with each axis comprising a histogram binning with 100 bins on each axis.

These means (red dots) from the 1000 element segment appear in the plot in figure 2 middle row, indicating the portion of the data from which they were taken. Note that the inverse relationship between LASY and CE is not apparent when plotting a subset of only 1000 elements, but when considering the entire data set the large scale trend becomes clearly evident. The blue dots in figure 2 middle indicate the means of all of the approximately 3000 individual 100 element segments from the entire data set. The density plot for the 300,000 individual *LASY, CE* pairs is shown in figure 2 bottom panel. For high values of *LASY* (low synchrony) the CE of the individual vertices are clustered in the region of small counts at the bottom of the vertical (*CE*) axis, whereas for low values of *LASY* (high synchrony) the density at the bottom becomes small as the trend toward larger values of *CE* is evident.

### The inverse relationship between *LASY* and *CE* is consistent across subjects and scans

*LASY* and *CE* were generated from the functional data of each of 21 subjects with six resting fMRI scans acquired during the same scan session for each subject. The CE measure was computed at correlation thresholds of *r* > 0.75, 0.60, 0.50 and 0.40. LOESS (local regression) curves were calculated from the data points generated for the *LASY,CE* plots as displayed in figure 2 middle row for all 126 scans at the four correlation threshold levels. The LOESS regressions were performed in Matlab (see ‘Statistics’ in method for the LOESS plot parameters used). Figure 3 presents the LOESS curves derived from the scans separately for each rest condition and at the correlation thresholds of 0.75 and 0.60. The results for the correlation thresholds *r* = 0.5 and 0.4 are presented in supplementary material figure 1. The LOESS curves clearly indicate that for all scans and correlation threshold levels the largest *CE* always occurs at low *LASY*.

**Figure 3.**
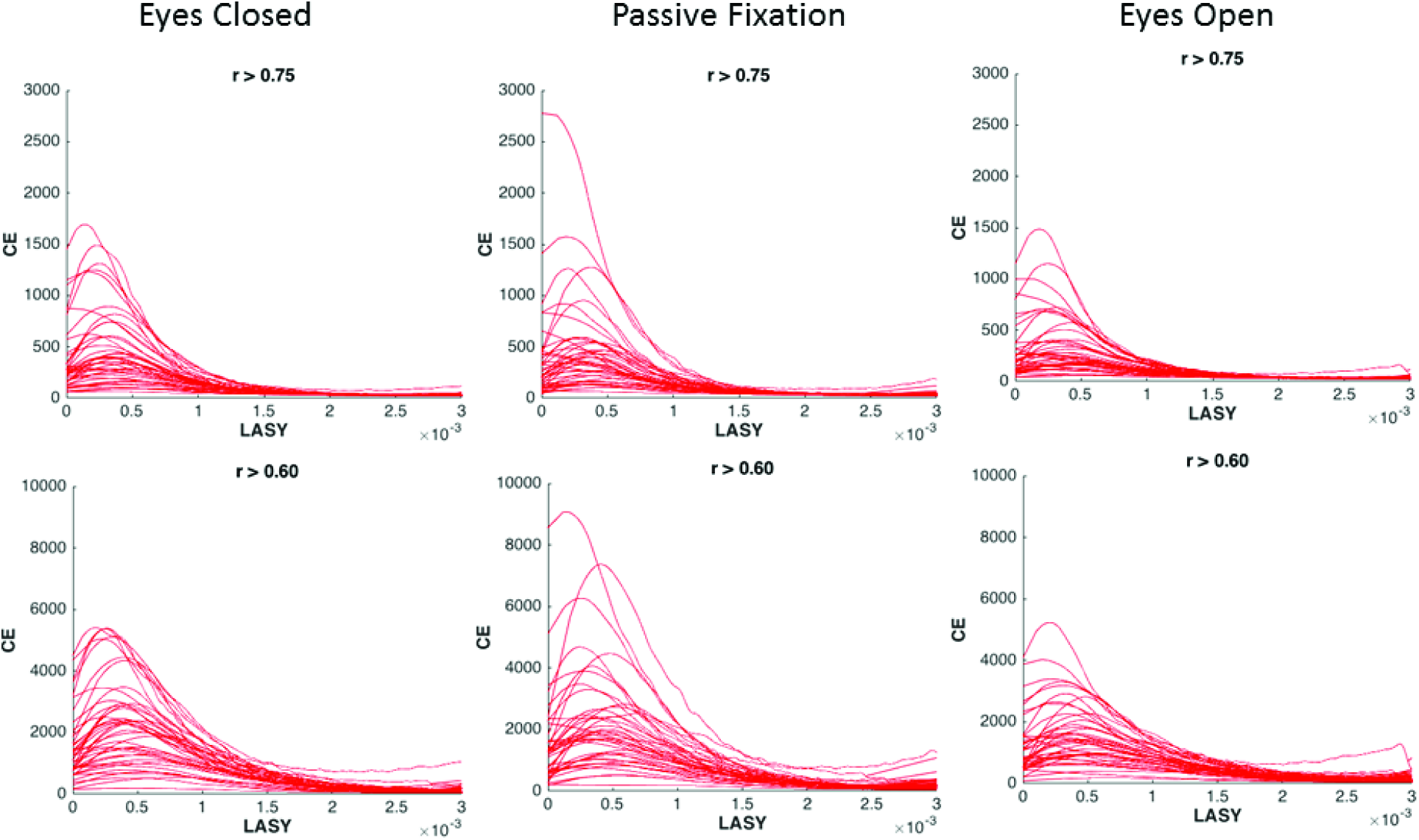
A consistent relationship across scans. LOESS (local regression) curves generated from the *LASY,CE* plots of all 126 scans at the correlation threshold values 0.75 and 0.60. The results from 42 scans (2 scans in each rest condition from each of 21 subjects) are presented in each plot. The largest *CE* (count of vertices in thresholded seed based connectivity maps) occur at vertices with highly synchronous local neighborhoods.

In order to quantify this relationship, we computed the Spearman’s rank order correlation between *LASY* and *CE* independently for each of the 126 scans and at each of the four correlation levels. While the data were averaged for visualization purposes, these tests were performed on the full set of ~300,000 *LASY, CE* pairs for each scan. For each of these (126 * 4) tests the estimated *p*-value was < 10e^−96^. All correlations were negative indicating the inverse relation between *LASY* and *CE*. The histograms of these correlation values for each correlation threshold are displayed in figure 4.

**Figure 4.**
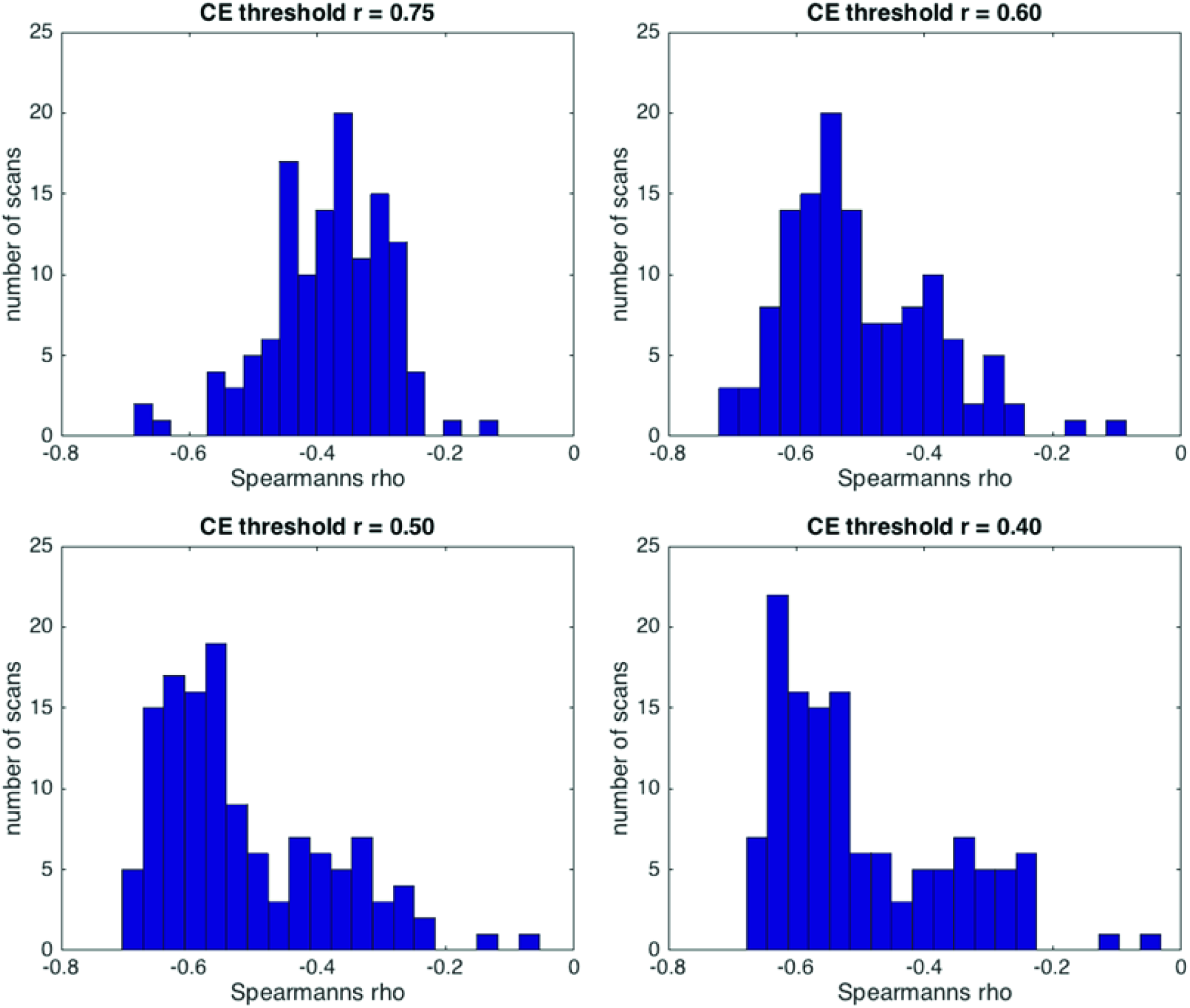
A strong inverse relationship between LASY and CE for all scans. Distribution of the Spearman’s linear correlation coefficient between *LASY* and *CE* tested independently for each of the 126 scans. This is presented for CE maps thresholded at *r* = 075, *r* = 60, *r* = 50 and *r* = 40. Note: all correlations are negative reflecting the inverse relationship between *LASY* and *CE*. The estimated p-value for all 4*126 tests was < 10e^−96^

### The inverse relationship between *LASY* and *CE* is dynamically preserved across repeated within session scans

In this section we demonstrate that *LASY* and *CE* are inversely correlated across time while controlling for the temporal SNR (*TSNR*) of the data. The previous section demonstrated that the inverse relationship between LASY and CE holds across vertices (space). Here, for each of the 21 subjects we analyze the relationship across the six repeated within session scans on an individual vertex level. We first fit a linear model *CE* = *β*_0_ + *β*_1_*LASY* + *β*_2_*TSNR* at each of the vertices where the dynamic range of LASY > 10^−4^. The key effect in which we are interested is captured by the properties of *β*_1_. For each of the 21 subjects, we take the mean of the estimated *β*_1_ across all the vertices. We also measure the percentage of *β*_1_s that are negative. The sign of the mean *β*_1_ and the proportion of the negative slopes indicate the type of relationship between *LASY* and *CE*. Fig. 5 shows that *LASY* and *CE* are inversely correlated. Fig. 5(a) shows the kernel density plots^28^ of the mean *β*_1_s using all the 21 subjects. Fig. 5(b) shows the density plots of the proportion of the negative *β*_1_s. We can observe that the mean slopes are *always* negative and the proportion of negative slopes is greater than 60% for all the different thresholds used in deriving *CE*. Here it is not essential to quantify the magnitude of the relationship but to show that it consistently indicates an inverse relationship between LASY and CE across time.

### 4.0 Discussion

Our results establish a consistent inverse relationship between local low temporal frequency fMRI signal synchrony (*LASY*) and the spatial extent of functional connectivity (CE) as measured by seed time series correlation based fcMRI. The LOESS plots of the relationship between *CE* and *LASY* (figure 3) indicate that vertices with high local synchrony are engaged in the largest and most highly correlated functional connectivity networks. The calculation of Spearman’s rho between *LASY* and *CE* confirm the inverse relationship (figure 4). The assessment of the relationship across repeated within session scans demonstrates that the relationship is also dynamically preserved across time(figure 5). We present in the supplementary material a detailed analysis of the influence of bulk motion, white matter average signal, cerebro-spinal fluid average signal and cardiac and respiratory induced motion. We demonstrate that the relationship is preserved after controlling for these influences. Especially for scans having high Spearmann’s rho after controlling for artifacts, the additional control measures are seen to increase the strength of the relationship. We also show that the correlations between *LASY* and a number of other possible confounding influences including temporal SNR, local curvature and vertex-wise area are also negligible in relation to the magnitude of the relation between *CE* and *LASY (supplementary material)*. Given the number of subjects and scans tested and the consistency of the relationship, we conclude that the result generalizes to healthy young subjects.

**Fig. 5.**
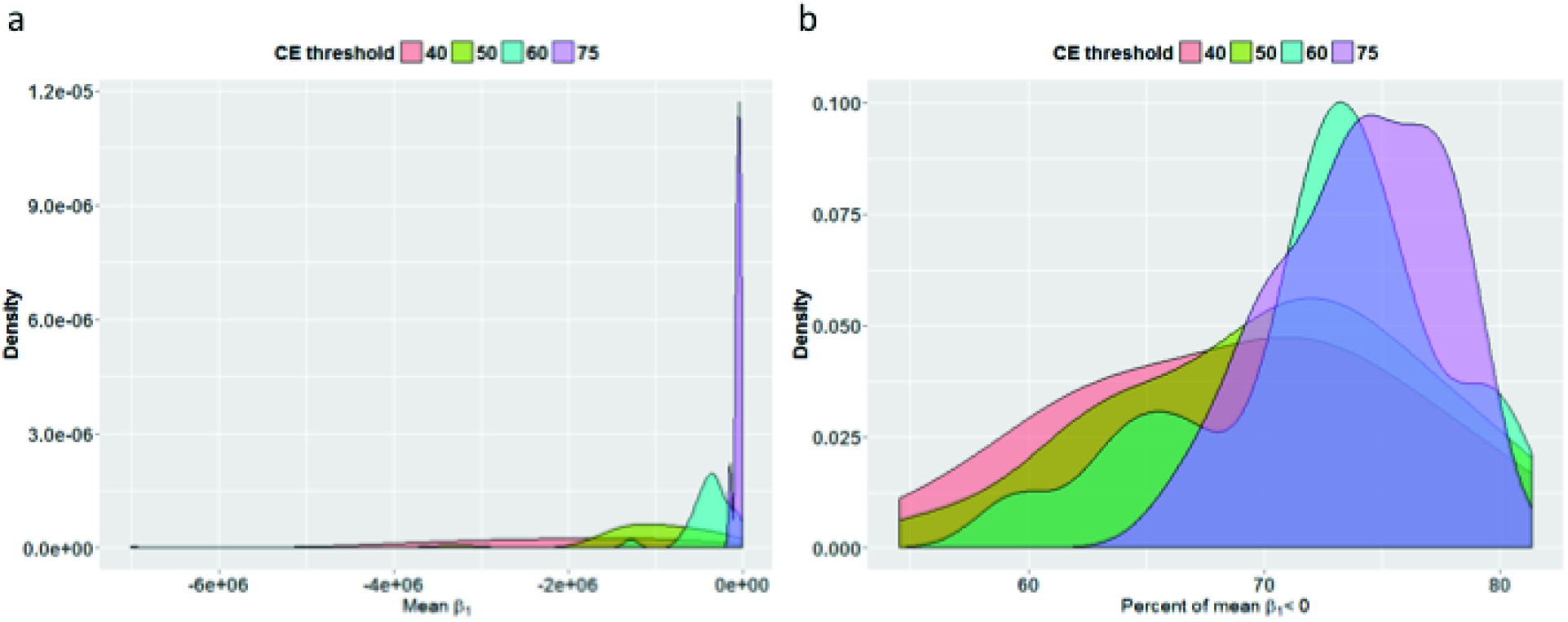
Relationship between LASY and CE is dynamically preserved across scans. The figures show that the inverse relationship between LASY and CE is dynamically preserved across repeated within session scans. Data at the individual vertex level are pooled across all 21 subjects. Kernel density plots of the (a) mean slopes (*β*_1_s) and (b) percentage of negative slopes for different thresholds used in deriving *CE*. The vertical axis is the density as the area under the curve is one.

One previous study showing evidence in support of a relationship between regional homogeneity (REHO) and global functional connectivity only demonstrated such a relationship for data averaged across subjects, and no individual subjects level data were presented^29^. This is not equivalent to the result we have presented, which is that *LASY* consistently shows a strong inverse relationship with seed-based global functional connectivity on an individual subject basis for each of the 126 scans tested.

There is always concern that the analysis may be affected by undetected and unaccounted for sources of artifact. We have however applied an extensive processing scheme to control for the influence of artifacts. We also demonstrate that the inverse relationship is consistent both before and after the artifact control scheme (supplementary material). Though there is a perception that group studies of resting state fMRI are less affected by “noise”, data averaging only provides an increase in SNR if the data represent repeated measurements of a single value with additive noise. There is no assumption in group studies of resting state fMRI that measurements of the correlation metric at the same location across subjects represent the same value, rather it is considered that the measurements represent a range of values of connectivity across subjects and the purpose of the analysis is to show the mean of one group is different from the mean of the second group.

Given the relationship demonstrated, the cortical surface with *LASY* overlain presents a map of the spatial variations in the scale of connectivity. These maps are essentially functional parcellations as a change in the color/intensity indicates a change in the scale of connectivity. The local asynchrony maps presented in figure 1 and supplementary material figures 2 and 3 indicate an intricate pattern of high connectivity vertices (vertices with low *LASY*), embedded in a cortex composed of much lower connectivity vertices.

The *LASY* maps do not depict functional connectivity that is smoothly varying or approximately locally homogenous. On the contrary, they point to a much more fine scale and irregular structure than the one represented by graph nodes and parcellations commonly defined in fcMRI studies^17, 18, 30, 31^. The spatial pattern of functional connectivity presented here seem to indicate that the functional architecture of the cortex may be better characterized by a collectivist model of many interacting structural elements than by a connectionist model of a relatively small number of spatio-temporally static nodes^32^. Differences in the spatial resolution of nodes in graph analysis of fcMRI connectomics result in large changes in graph theoretic measures^27^. One study indicated that the most serious confound for accurate network measure estimation was inaccurate functional node specification^33^. The requirement for a node to have homogenous functional connectivity would imply that regions defined as nodes be constant in the *LASY measure* to some level of approximation. Simply put, if a node contains both a high synchrony and low synchrony area then the connectivity is not homogenous within the node. Since the local synchrony only predicts the scale of connectivity and not the actual connectivity pattern it would be possible that within a node that is homogenous in *LASY* the spatial pattern of connectivity changes but the size of networks remains the same. This implies that the requirement for constant *LASY* is a necessary but not sufficient condition for functional homogeneity of the node.

There is a seeming contradiction in resting state networks research. On the one hand, a fairly large number of test-retest reliability results has been published^34–36^. On the other hand there a growing literature describing changes in network configurations related to tasks^37, 38^ and also a growing literature devoted to dynamic functional connectivity^39–41^. The networks cannot be both highly replicable and also highly dynamic. This inconsistency is possibly obscured by the practice of averaging across subjects and space. However, it is a simple statistical fact that the average of a population quantity may in fact represent none of the individual measurements. On reflection the true nature of the spatial patterns of connectivity can in the end only be resolved by an analysis of the patterns of connectivity expressed by individual subjects^42^ and the degree of temporal dynamics manifest at the individual subject level. The topic of inter-individual differences in functional architecture has been little explored^43^ and may be especially important in the area of psychiatric disorders^44^. Though the issue of poor replication of neuroimaging results has received a great deal of attention, questions related to the lack of anatomical accuracy and inter individual differences have been largely avoided^45^. In light of these issues the individual level connectivity analysis must be performed without the bias of prior spatial models of connectivity patterns. The main contribution of the present work is to provide an analysis methodology that indicates changes in connectivity across the entire cortex at the imaging voxel resolution and on an individual subject basis. This analysis is data driven and requires no prior spatial model of connectivity.

## Material and Methods

### Participants and fMRI Data Acquisition

Written informed consent was obtained from subjects prior to each scanning session in accordance with an approved University of Wisconsin-Madison IRB protocol. Twenty-one healthy adults (mean age 23.6, 12 Female) with no prior history of neurological or psychological disorders were studied. Six resting fMRI scans and a T1-weighted image were acquired from each subject during a single acquisition session. All scans were acquired using a 3 T GE scanner (MR750, General Electric Healthcare, Waukesha, WI) using the product 8-channel receive-only radio frequency (RF) coil. Each fMRI scan was 10 minutes in length and acquired with the same echo planar imaging (EPI) sequence (TR = 2.6 s, TE = 25 ms, flip angle = 60 degrees, FOV = 224 mm × 224 mm, matrix size = 64 × 64, slice thickness = 3.5 mm, number of slices = 40). Two eyes open resting conditions, two eyes closed resting conditions, and two passive fixation scans were acquired from each subject during one scan session. The participants were instructed to relax and lie still in the scanner while remaining “calm, still, and awake”. For the passive fixation scans subjects were instructed to keep their eyes open with their gaze fixated on a cross back-projected onto a screen via an LCD projector (Avotec, Inc., Stuart, FL). Subjects were instructed to keep their eyes open/closed for the EO/EC conditions and were allowed to blink if necessary. T1-weighted structural images were acquired before the functional images using a 3D BRAVO inversion-recovery prepped fast gradient echo sequence with the following parameters: TR = 8.13 ms, TE = 3.18 ms, TI = 450 ms, flip angle = 12 degrees, FOV = 256 mm x 256 mm, matrix size = 256 × 256, slice thickness = 1 mm, number of slices = 156.

### Preprocessing, Registration and Resampling

All functional images were preprocessed by performing motion correction, slice timing correction and band pass filtering [0.01, 0.1] Hz using AFNI ^46^. Two sets of functional data were prepared for comparison. A first set was prepared by performing in addition to the previously stated preprocessing a stringent motion censoring rejecting all time points with more than 0.25 mm motion, measured from the Euclidean norm of the frame-to-frame difference in the 6 estimated realignment parameters^47, 48^.

The second set of functional data was prepared by first reducing the influence of physiological noise using RETROICOR^49^ between the motion correction and slice timing correction steps^50^, and then regressing out the average white matter (WM) signal (over an eroded white matter mask), the average CSF signal (over an eroded CSF mask), the derivatives of the average WM and CSF signals, and the 6 realignment parameters. All scans used in the study had high quality cardiac and respiratory signals. Time points where frame-to-frame motion exceeded 0.25 mm were censored and not included in this nuisance regression.

Surface reconstruction was performed on each subjects T1 weighted image using Freesurfer ^51^. The functional MRI data were registered to the subjects cortical surface using boundary-based linear registration ^52^ and that registration was used to resample each functional time point volume onto the cortical surface using nearest neighbor interpolation. The fMRI data were sampled from the midpoint between the pial and grey/white surface, thus minimizing the partial volume averaging of cerebrospinal fluid (CSF) and white matter. The cortical fMRI data were reordered to produce a matrix with one column for each vertex on both hemispheres of the cortical surface containing the functional time series resampled to the vertices location. Finally, each time series was normalized to zero mean and unit variance. The number of vertices comprising both hemispheres of an individual subject’s cortical reconstruction averages roughly 300,000 and varies by +/− 30,000 due to variations in brain size and cortical folding pattern. The area of the medial wall of the Freesurfer cortical surfaces, which is not cortex and is a computational convenience of the reconstruction, was not included in any analyses.

### Local asynchrony calculation

The local asynchrony (*LASY*) is computed as outlined in Blumensath et al. ^17^, and was termed stability map in that study. To calculate *LASY* at a vertex *v* we first construct the set of spatially connected vertices on the cortical surface within 3 mm Euclidean distance of *v* by a region growing procedure (additional details provided in the supporting material). The region growing procedure used prevents cortex which is within 3 mm Euclidian distance of *v* but significantly more distant in geodesic distance on the surface from being included. Such cases arise for example when sulci are very thin and opposite banks of sulci are very close. Euclidean distance in the volume rather than geodesic distance on the surface was used due to the extensive computation required to produce approximate geodesic distance measures.

The number of vertices within 3 mm of a selected vertex will vary as it depends on the folding pattern of the cortex near *v*. The surface area of cortex represented by a single vertex varies slightly but the distribution of area of a vertex is sharply peaked at a value slightly smaller than 1 mm^2^. The choice of 3 mm radius for the neighborhood calculation was motivated by the largest EPI voxel spatial dimension, which is 3.5 mm. We need to sample from at least 2 functional time series in each dimension when calculating *LASY (synchrony cannot be estimated by sampling a single time series)*. We also wished to obtain the highest possible resolution, which motivated not choosing a larger and thus lower resolution/higher radius neighborhood. The 3 mm radius is the same as used in the parcellation procedure from which the implementation of the algorithm was taken ^17^. We remind the reader that the functional data were resampled using nearest neighborhood interpolation which introduces no smoothing and no additional smoothing was performed.

Having obtained the list of vertices to be included in the neighborhood, we construct an *M*x*N* matrix where *M* is the number of time points in the fMRI time series and *N* is the number of vertices within the 3 mm neighborhood of *v*, including *v*.

We calculate *LASY* at *v* as the mean square error between all time-series in the neighborhood of *v* and the mean time series in the neighborhood of *v*. In the methods section we reserve the lower case lasy for a single value and upper case LASY for a vector of values.

For each vertex *v*, *lasyv* is thus the scalar corresponding to the variance of the set of numbers in the entire matrix *Av*, not the vector of row-wise or column-wise variances.

We generate the vector *LASY =* [*lasyv*] of local asynchrony measures at each vertex on both hemispheres of the cortical surface.

Note that while the local asynchrony, *lasy*, is derived by computing the variance within a local neighborhood, it provides a measure of synchrony because the mean signal across vertices within the local neighborhood at each time point has been subtracted prior to computing the variance. An example matrix *D* consisting of *N* copies of the same time series in the columns, that is perfectly synchronous data, would result in all entries of *A* be equal to 0 and *lasyv* = 0. Also time points where all time series values are equal would result in a zero row in the *A* matrix and contribute nothing to *lasyv*. NOTE: *lasy* = 0 indicates perfect synchrony and larger *lasy* values indicate lower synchrony. This asynchrony calculation and the prerequisite neighborhood information were generated using the Matlab code implemented for a functional parcellation procedure published earlier ^17^. The measure we term *lasy* was used as a regional homogeneity measure in the initial computational steps of the parcellation algorithm published by Blumensath and colleagues. The remaining steps of the parcellation scheme are unrelated to the analysis presented here.

### Relationship between global connectivity and local synchrony

In order to investigate the relationship between local synchrony and global functional connectivity we first generate fcMRI correlation maps (statistical parametric maps) for each vertex on the cortical surface. Specifically, we compute the vector *cv* = corr(*X,seed_v_*). Here *X* is the matrix of time series across all vertices of both hemispheres of the cortical surface and is on the order of 300 time points * 300,000 vertices for our data. The exact number of time points in the matrix for a scan will vary somewhat from one subject to another as the motion censoring procedure may have removed some time points. Note that the time series in the matrix *X* are not normalized to unit energy and zero mean as are the data used to calculate *lasy*. The vector *seed*v contains the fMRI time series at the vertex *v* and is taken from the matrix *X*; corr is the Pearson linear correlation coefficient.

The number of vertices with correlation coefficients in *cv* above a specified threshold *r* is defined to be the **connectivity extent** for the vertex *v*, or c*er,v* and *CEr* is the vector of values of c*<u>er,v</u>* for all vertices across both hemispheres of the cortical surface, excluding the medial wall.

At this point in the procedure the ordering of the values comprising *LASY* and *CEr* is that of vertices on the cortical surface as implemented internally in the Freesurfer software.

In determining the relationship between two variables it is desirable to have the independent variable ordered in increasing order. To achieve this, we sort *LASY* in ascending order.

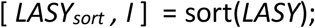

where *I* is the reordering. In order to maintain the correspondence between *lasyv* and *cer,v* pairs we apply the reordering to the correlation count vector to get *CEsortI* = reorder(*CEr,I*), where we have for brevity omitted the subscript indicating the correlation threshold *r*.

We now have obtained *F* = (*CE_sortI_,LASY_sort_*), the mathematical relationship between the size of the thresholded connectivity maps and local synchrony. The calculation of *CE_sortI_* and *LASY_sort_* was performed for all 126 scans at each of four correlation thresholds, 0.75, 0.60, 0.50 and 0.40 used for thresholding the CE.

### Statistics

The LOESS plots were generated to provide a graphical representation of the relationship between LASY and CE using locally weighted linear regression to smooth the data. The LOESS method was used to fit a quadratic model using a window size of 5% of the range of LASY.

The Spearman’s rank order correlation or Spearman’s rho was used to quantitatively assess the relationship between LASY and CE. It is a nonparametric measure of rank correlation that assesses how well the relationship between two variables can be described using a monotonic function.

General linear models were used to establish that the inverse relationship between LASY and CE was dynamically preserved across repeated within session scans. Kernel density estimates of the distributions of the mean as well as the percentage of negative model coefficients were used to provide graphical representation of this inverse relationship.

## Data and Code availability

The code used to perform these calculations is available for download, as well as a subset of the data used for this study (NITRC fcon 1000 (CoRR) website, http://fcon_1000.projects.nitrc.org/indi/CoRR/html/).

## Acknowledgments

Center For High Throughput Computing (CHTC) in the Department of Computer Sciences. The CHTC is supported by UW-Madison, the Wisconsin Alumni Research Foundation, the Wisconsin Institutes for Discovery, the National Science Foundation, and the Advanced Computing Infrastructure, and is an active member of the Open Science Grid, which is supported by the National Science Foundation and the U.S. Department of Energy’s Office of Science. This research was supported by NIH Grant RC1MH090912.

## Author contributions

G.K. Developed the methods, performed the analysis and wrote the manuscript. R.B. performed the artifact control regressions. A.A. wrote the manuscript and guided the analysis. D.D. wrote the paper and guided the analysis. N.A. performed the analysis of the temporal structure of the relationship. T.B. implemented and provided software for the implementation of the synchrony measure. M.F.C. wrote the paper and guided the analysis. B.T. L.M. implemented the large scale computing methods used.

